# Non-coding circular RNAs repertoire and expression profile during *Brassica rapa* pollen development

**DOI:** 10.1101/2021.07.29.454397

**Authors:** Saeid Babaei, Mohan B. Singh, Prem L Bhalla

**Affiliations:** Plant Molecular Biology and Biotechnology Laboratory, Faculty of Veterinary and Agricultural Sciences, The University of Melbourne, Parkville, Melbourne, VIC 3010, Australia

**Keywords:** *Brassica rapa*, circRNAs, pollen development, RNA sequencing

## Abstract

Circular RNAs (circRNAs) are covalently closed long non-coding RNA (lncRNA) molecules generated by the back-splicing of exons from linear precursor mRNAs. Though linear lncRNAs have been shown to play important regulatory roles in diverse biological and developmental processes, little is known about the role of their circular counterparts. In this study, we have performed high-throughput RNA sequencing to delineate the expression profile and potential function of circRNAs during the five stages of pollen development in *Brassica rapa*. A total of 1180 circRNAs were detected in pollen development, of which 367 showed stage-specific expression patterns. Functional enrichment and metabolic pathway analysis showed that the parent genes of circRNAs were mainly involved in pollen-related molecular and biological processes such as mitotic and meiosis cell division, DNA processes, protein synthesis, protein modification, and polysaccharide biosynthesis. Moreover, by predicting the circRNA-miRNA network from our differentially expressed circRNAs, we found 88 circRNAs with potential miRNA binding sites suggesting their role in post-transcriptional regulation of the genes. Finally, we confirmed the back-splicing sites of 9 randomly selected circRNAS using divergent primers and Sanger sequencing. Our study presents the first systematic analysis of circular RNAs during pollen development and forms the basis of future studies for unlocking complex gene regulatory networks underpinning reproduction in flowering plants.

## Introduction

Transcriptome-wide sequencing studies have shown that the genomic information is extensively transcribed into protein-coding RNAs and RNAs with no protein coding potential but play widely important regulatory roles. Although alternative splicing and post-transcriptional events can generate numerous mRNA isoforms to increase the protein-coding and regulatory capacity of the eukaryotic genomes, these RNAs comprise only a small percentage (usually about 1-2%) of all RNAs in the cell (de Klerk and AC‘t Hoen 2015). The majority of expressed RNAs are non-coding RNAs based on their size and structure divided into multiple categories such as small non-coding RNAs, lncRNAs, and circRNAs (Mattick and Makunin 2006; Cech and Steitz 2014; Meng et al. 2017). Non-coding RNAs have been known to be key regulatory factors controlling gene expression at the post-transcriptional, transcriptional, and epigenetic levels (Kaikkonen et al. 2011). Among non-coding RNAs, circRNAs are covalently closed single-stranded RNA molecules (with no 5’ caps or 3’ poly-A tails) following transcription from a wide range of genomic positions, including exons, introns, and intergenic regions, undergo a non-canonical splicing event termed as back-splicing. (Zhou et al. 2020). Despite being known for decades, circRNAs have been considered as aberrant byproducts of mis-splicing events (Guria et al. 2020). With the advances in RNA-seq technology and bioinformatics algorithms, thousands of circRNAs have been identified in all eukaryotic tree of life and revealed them to have cell and tissue-specific expression patterns (Wang et al. 2014; Zhang et al. 2020b). Earlier studies have demonstrated that lncRNAs play important roles in reproductive-related processes such as floral transition and flower development (Golicz et al. 2018). Cold-Assisted Intronic Noncoding RNA (*COLDAIR*) and Cold-Induced Long Antisense Intragenic RNA (*COOLAIR*) are lncRNAs that control flowering in *Arabidopsis* by repressing Flowering Locus C (*FLC*) through epigenetic mechanisms (Heo and Sung 2011; Csorba et al. 2014). In rice, it has been revealed that more than 700 long intergenic non-coding RNAs are essential factors for biogenesis of phased small interfering RNAs that associate with germline-specific Argonaute protein MEL1, which itself is involved in the development of pre-meiotic germ cells, suggesting that rice lncRNAs are important elements in reproduction (Komiya et al. 2014).

Cell-specific expression of thousands of protein-coding transcripts underpins pollen developmental processes (Singh and Bhalla 2007; Haerizadeh et al. 2009; Russell et al. 2012). Cis-regulatory elements, tissue-specific promoters, histone modifications, and DNA methylation have also been reported to be involved in the regulatory network of pollen development (Janousek et al. 2000; Sharma et al. 2011; Okada et al. 2006; Singh et al. 2003; Xu et al. 1999). lncRNAs have also been implicated in controlling pollen reproductive developmental processes (Golicz et al. 2018). For example, Long-Day Specific Male Fertility Associated RNA (*LDMAR*) and *Zm401* are lncRNAs required for pollen development in rice and maize, respectively (Ma et al. 2008; Ding et al. 2012). A recent study in *Brassica rapa* found that more than 12000 lncRNAs were expressed during pollen development and fertilization (Huang et al. 2018). In addition, recent studies have implicated circRNAs in the regulation of pollen development. For instance, in rice, 186 differentially expressed circRNAs have been identified in different pollen developmental stages during the fertility transition of the photo-thermosensitive genic male sterile line (Wang et al. 2019c). In soybean, 2867 circRNAs were identified in flower buds of the cytoplasmic male sterile line (NJCMS1A) and its maintainer (NJCMS1B), of which 1009 circRNAs were differentially expressed between two lines pointing towards the role of circRNAs in flower and pollen development (Chen et al. 2018). Similar work in *B. campestris*, revealed 31 differentially expressed circRNAs between cytoplasm male sterile and fertile lines involved in anther development (Liang et al. 2019). To further improve our understanding of the role of circRNAs in pollen development, we performed a time series of RNA-seq experiments during pollen development in *B. rapa* (AA, 2n = 20)., Using three different bioinformatic prediction tools, we identified 1180 circRNAs expressed in five pollen developmental stages. We performed differential gene expression analysis, functional annotation, and pathway enrichment analysis to explore the potential function of circRNAs during pollen development. We also predicted miRNA-circRNA interactions to investigate the potential role of circRNAs as competing endogenous RNAs (ceRNAs) in post-transcriptional gene regulation.

## Materials and Methods

### Plant materials, growth conditions, and sample collection

In this study, *Brassica rapa* accession no. ATC 92270 Y.S (AND)-168 were grown at 21/18°C day/night under 16/8 hours’ light/dark (200μmol m^−2^s^−1^ light intensity) with 60% humidity. Based on the size of floral buds, five pollen developmental stages were identified: pollen mother cells: ≤1 mm buds, tetrad: 1.2-2 mm buds, uninucleate pollen: 2-2.6 mm buds, binucleate pollens: 3-4 mm buds, and mature pollen: 5-6 mm buds (Huang et al. 2018) (Figure 1). Collected buds were immediately transferred to a petri dish containing modified 0.5 X B5 medium (13% sucrose) on ice. Pollen from the buds in their binucleate and mature pollen stages was released by dissecting out the anthers and squashing them in a modified B5 medium. Because the buds in the first three groups (pollen mother cells, tetrad, and uninucleate pollen) were too small, the whole buds were squashed in the modified B5 medium to release the pollens. The resulting suspensions were then filtered using a 40μm mesh (pluriStrainer Mini 40μm, pluriSelect, Germany) into 2ml tubes. Tubes were then centrifuged at 4°C for 3 minutes at 150g, and the supernatant was discarded. The pellet was washed with a modified 0.5 X B5 medium, and finally, pure pollen grains were collected from the medium using centrifugation at 4°C for 3 minutes at 150g. After discarding the medium, the pellet was immediately frozen in liquid nitrogen and stored at −80°C for later use (Lohani et al. 2020).

**Figure 1.**
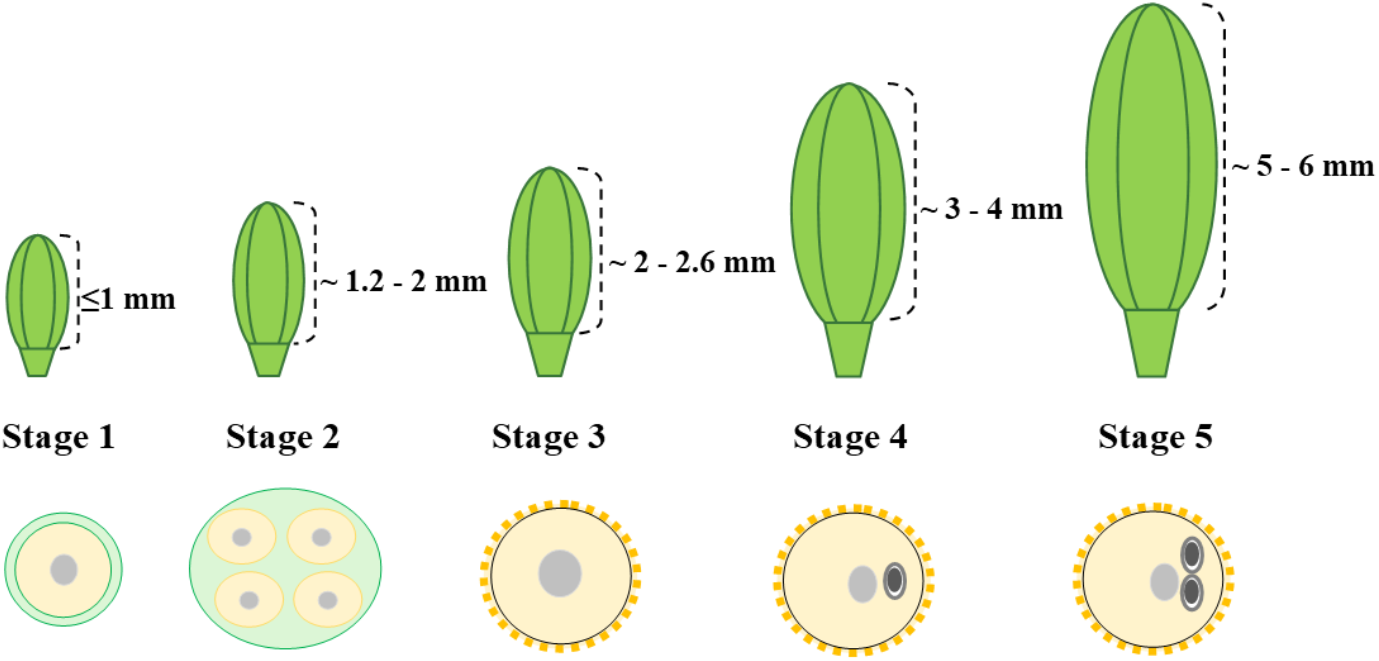
A schematic representation of floral buds at different developmental stages and their corresponding microspores. Stage 1: pollen mother cells, stage 2: tetrad, stage 3: uninucleate pollen, stage 4: binucleate pollen, and stage 5: mature pollen.

#### RNA extraction and sequencing

The total RNA was isolated from collected pollen samples using the mirVana™ miRNA Isolation Kit (Thermo-Fisher; Part Numbers AM1560, AM1561) according to the manufacturer’s instructions. To remove contaminated DNA, all the isolated RNA samples were treated with TURBO™ DNase (Ambion) and then submitted for sequencing. The RNA-seq libraries were then constructed using Illumina TruSeq stranded kit following the rRNA depletion step, and the single-end reads (100 bp) sequencing was done at the Australian Genome Research Facility (AGRF), Melbourne.

#### Identification and Differential Expression of Circular RNAs

After obtaining clean reads by removing low-quality and adaptor sequences, reads were mapped to the *B. rapa* reference genome (v3.0, http://brassicadb.org/brad/) (Wang et al. 2011) using Bowtie2 (v2.3.5.1) (Langmead and Salzberg 2012), STAR (v2.7.5a) (Dobin et al. 2013), and BWA (v0.7.17, mem-T 19) (Li and Durbin 2009). Then, circRNAs were identified using three different tools, find_circ (v1.2) (Memczak et al. 2013), CIRCexplorer2 (v2.3.8) (Zhang et al. 2016), and CIRI2 (v2.0.6) (Gao et al. 2018) with their default parameters. The R-Bioconductor package Noiseq (v2.28.0) (Tarazona et al. 2011) (with norm= “tmm” and q=0.8) was used for quantification of expression level and detection of differentially expressed circRNAs between samples. Plots were generated using the R (v4.0.4) (Team 2013) with ggplot2 (v3.3.3) library (Ginestet 2011).

#### Functional Enrichment Analysis

Gene Ontology (GO) enrichment analysis was undertaken for functional categorization of the parental genes of differential expressed circRNAs by topGO package (v2.42.0) (Alexa and Rahnenfuhrer 2010), and KEGG (Kyoto Encyclopedia of Genes and Genomes) pathway enrichment analysis was conducted by submitting the sequence of differentially expressed circRNAs in KOBAS database (v3.0, http://kobas.cbi.pku.edu.cn/kobas3) (Xie et al. 2011).

#### Prediction of potential circRNA-miRNA interactions and visualization

The potential interaction between known miRNAs and our identified differential expressed circRNAs was carried out using web tool psRNATarget (v2, http://plantgrn.noble.org/psRNATarget/home) (Dai and Zhao 2011) with default parameters, except we set the Expectation value to 3.0. We selected “*Brassica rapa*, 157 published miRNA” as input miRNAs and the sequence of our identified differentially expressed circRNAs as the target sequences. Finally, the circRNA-miRNA network was visualized using Cytoscape software (v3.8.2) (Shannon et al. 2003).

#### Circular RNA validation

To validate our identified circRNAs that were predicted by bioinformatic approaches, PCR with divergent primers followed by Sanger sequencing was used. Briefly, total RNA from all five pollen developmental stages was reverse transcribed into complementary DNA (cDNA) using SuperScript™ III Reverse Transcriptase (Invitrogen) and random hexamers according to the manufacturer’s protocol. The sequences of 15 *in-silico* predicted circRNAs that were predicted to be expressed in developing pollen were used to design 15 pairs of divergent primers using Primer3web (v4.1.0) (Untergasser et al. 2012). The PCR procedure was as follows: initial step at 94°C for 3 min; followed by 40 cycles at 94°C for 30 s, appropriate annealing temperature (Table 1) for 45 s, and 72°C for 30 s; and then 1 cycle at 72°C for 7 min. The PCR products were then visualized by agarose gel (1.5%) electrophoresis, and the desired bands were recovered from the gel using Wizard® SV Gel and PCR Clean-Up System (Promega). Finally, the PCR products were cloned into pJET1.2/blunt vector using CloneJET PCR Cloning Kit (Thermo Scientific) and subjected to Sanger sequencing to confirm the back-spliced junction sites.

**Table 1.**
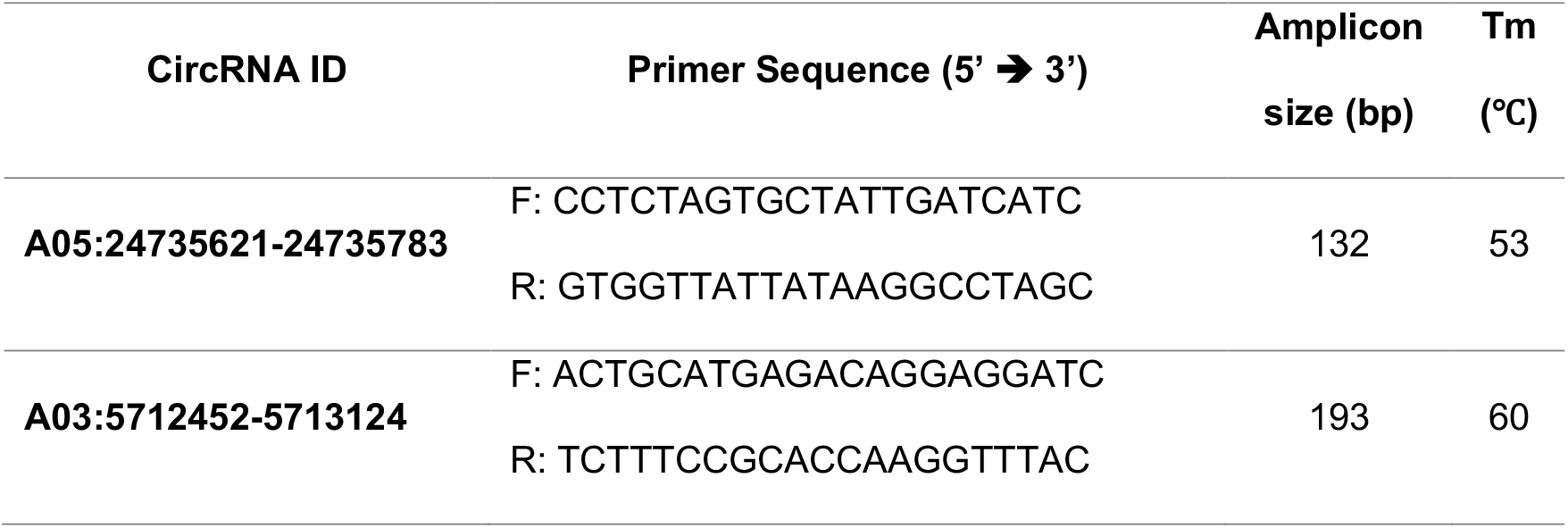
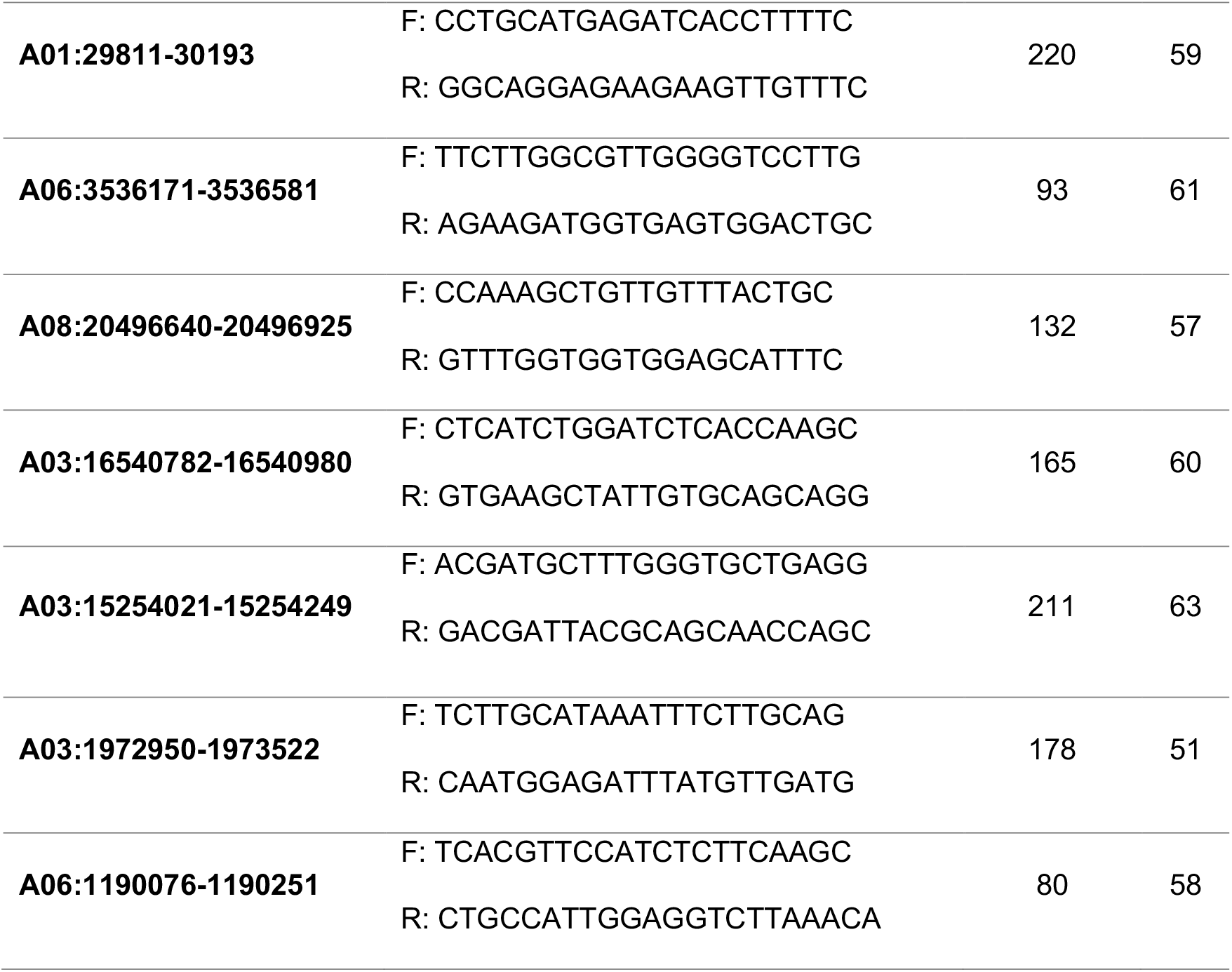
A list of divergent primers for circRNA conjunction validation.

## Results

### Identification and characterization of circRNAs

To explore the involvement of circRNAs in *B. rapa* pollen development, rRNA-depleted RNA libraries for five pollen developmental stages (pollen mother cells, tetrad, uninucleate pollen, binucleate pollen, and mature pollen) were constructed. A total of about 23 gigabytes of high-quality clean data with more than 233 million reads were obtained (Table 2). After analyzing the sequencing data with three circRNA identification tools, more than 2000 circRNAs were obtained (TableS1). Out of the total number of circRNAs, we identified 1180 unique circRNAs in all five pollen developmental stages with the portion of 824 circRNAs for Find_Circ, 380 circRNAs for CIRI2, and 137 circRNAs for CIRCExplorer2 (Figure 2, and TableS2). When comparing the number of circRNAs found by all three tools, only 6 circRNAs were common between all, and more circRNAs were common between CIRI2 and Find_Circ (Figure 2). Among 1180 identified circRNAs, 1034 (87.63%) circRNAs were predicted to have at least one exon from protein-coding genes (Figure 3A), 85 (7.20%) circRNAs, were generated from intergenic regions, and the remaining circRNAs, 61 (5.17%) were intronic. While 133 genes could produce two or more types of circRNAs through alternative back-splicing, 737 genes produced only one circRNA isoform (Figure 3B). Moreover, the parent genes of circRNAs were unevenly distributed on different chromosomes; chromosome nine with 184 circRNAs and chromosome four with 63 circRNAs produced the most and the least circRNAs, respectively, and 21 circRNAs were produced from four Scaffolds (Figure 3C). Finally, the length distribution of circRNAs had mainly distributed between 100 to 600 bp (Figure 3D).

**Table 2.** A summary statistics of sequencing data.

| **Libraries** | **No. of reads** | **GC content** | **Q20** | **Q30** |
| --- | --- | --- | --- | --- |
| **Pollen mother cells** | 45,634,748 | 45 | 97.25 | 94.84 |
| **Tetrad** | 49,878,644 | 45 | 97.49 | 95.20 |
| **Uninucleate pollen** | 44,187,256 | 45 | 94.22 | 90.65 |
| **Binucleate pollen** | 45,029,845 | 45 | 96.03 | 92.99 |
| **Mature pollen** | 48,755,777 | 45 | 97.21 | 94.75 |

**Figure 2.**
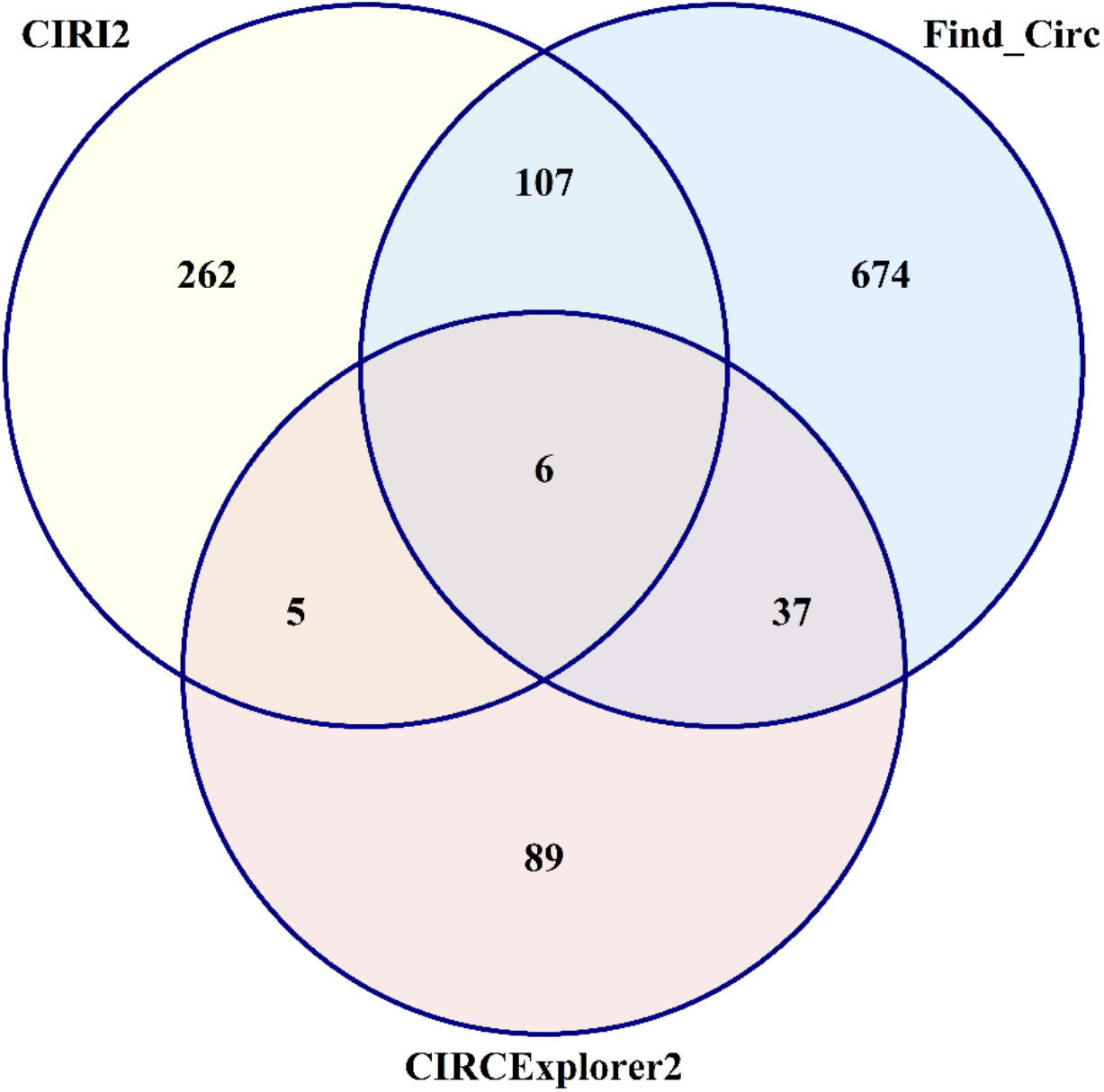
A Venn diagram of unique and shared circRNAs predicted by three algorithms: CIRCexplorer2, find_circ, and CIRI2.

**Figure 3.**
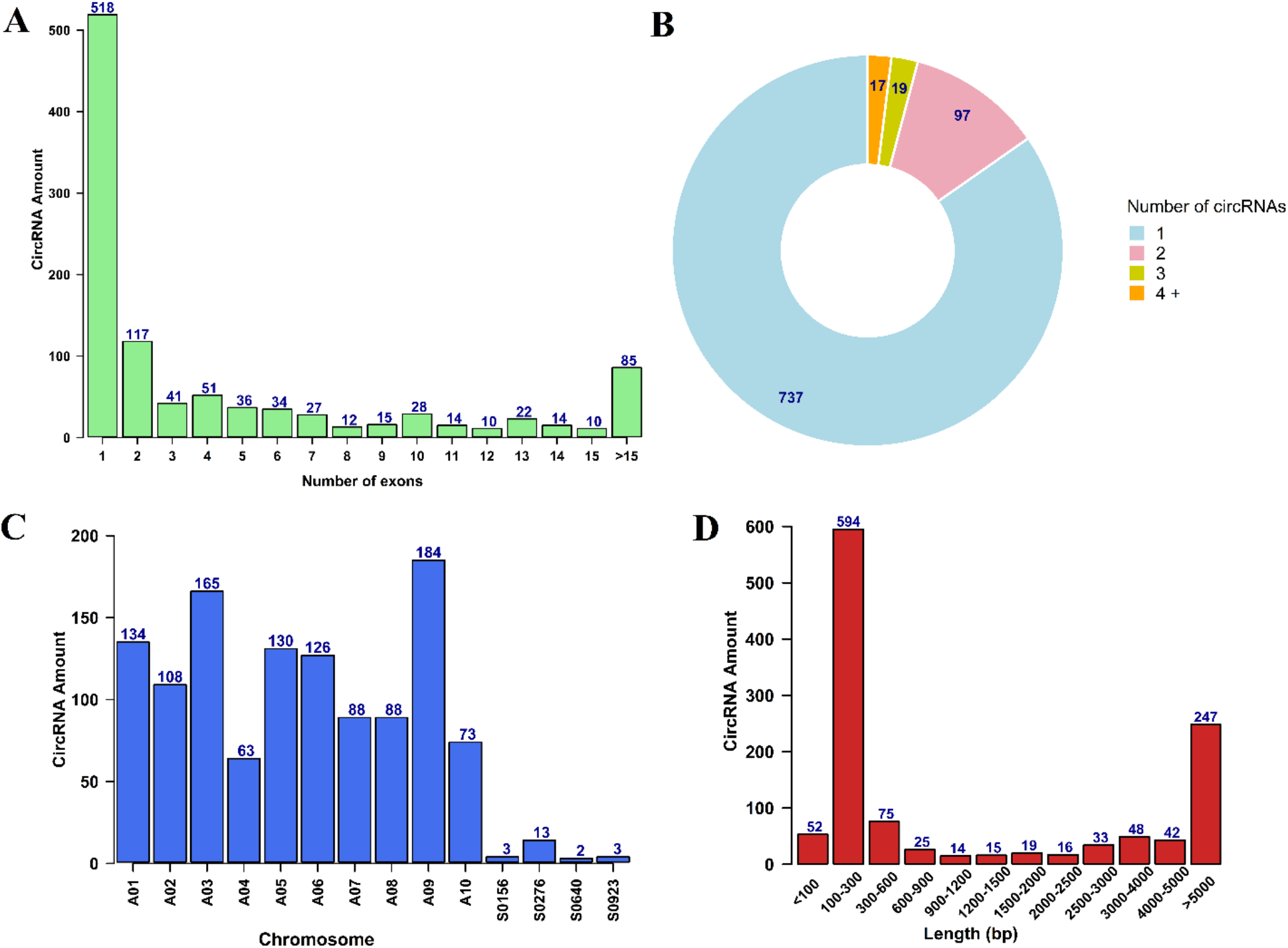
Genomic features of identified circRNAs during pollen development in *B. rapa*. (A) Distribution of the exon number per detected circRNA. (B) Distribution of the number of circRNA reads generated from the same parental gene. (C) The number of circRNAs detected in each chromosome. (D) Distribution of the length of circRNAs.

### The expression patterns of circRNAs in B. rapa pollen development

To investigate whether the circRNAs are expressed in a specific manner during *B. rapa* pollen development, we compared the expression patterns of circRNAs in each pollen developmental stage with its previous stage (TableS3). The results showed that 966 circRNAs had a significant differential expression, of which 367 circRNAs displayed stage-specific expression patterns.

Comparing the expression of circRNAs between stages, a more distinct expression was observed when pollen develops from binucleate pollen to mature pollen or develops from pollen mother cell to tetrad. (Figure 4A). Besides, more circRNAs tend to up-regulate during pollen development except when pollen develop from uninucleate pollen to binucleate pollen, where more circRNAs were down-regulated (Figure 4B). Finally, we visualized the expression profile of the 60 most significant (α ≥ 0.001) differential expressed circRNAs (TableS4) among different stages of pollen development using a heatmap (Figure 5). The expression analysis results indicated that circRNAs have a distinctive expression pattern during pollen development, pointing towards their defined stage-specific roles in the pollen developmental progression.

**Figure 4.**
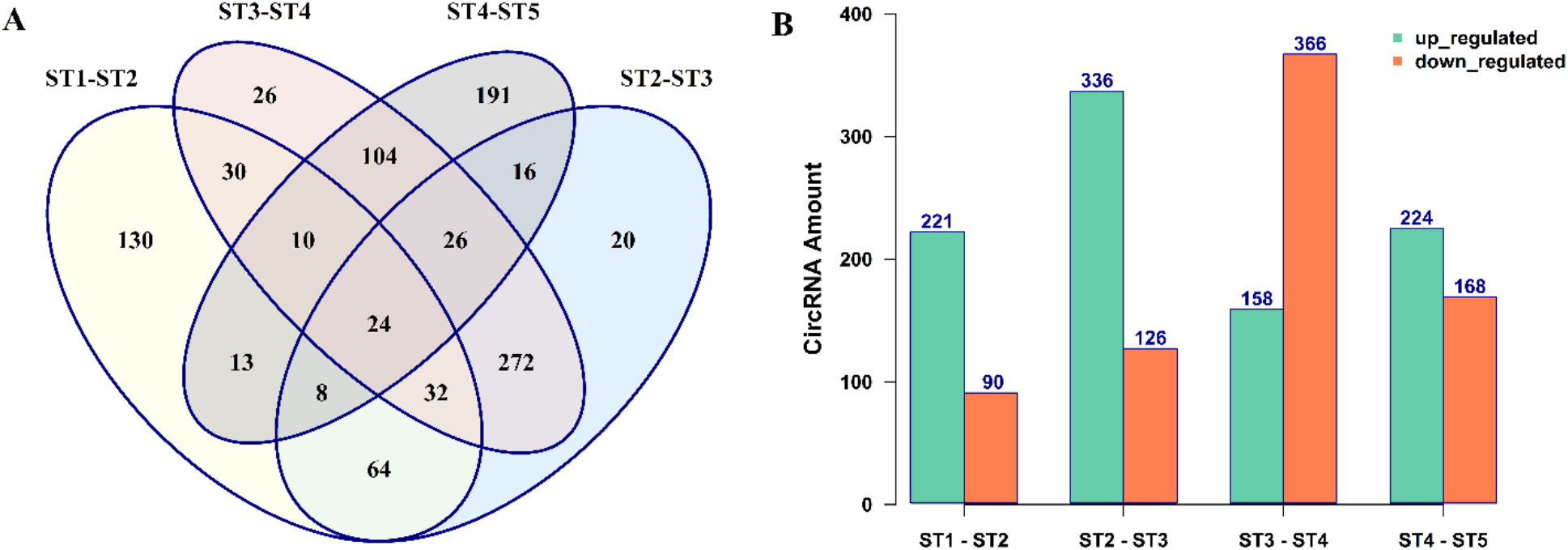
Differential expression patterns of circRNAs during pollen development in *B. rapa*. (A) A Venn diagram showing the number of differentially expressed circRNAs during pollen developmental stages. (B) Histograms represent the regulation of differentially expressed circRNAs during pollen developmental stages. ST1: stage 1 (pollen mother cells), ST2: stage 2 (Tetrad), ST3: stage 3 (Uninucleate pollen), ST4: stage 4 (Binucleate pollen), and ST5: stage 5 (Mature pollen).

**Figure 5.**
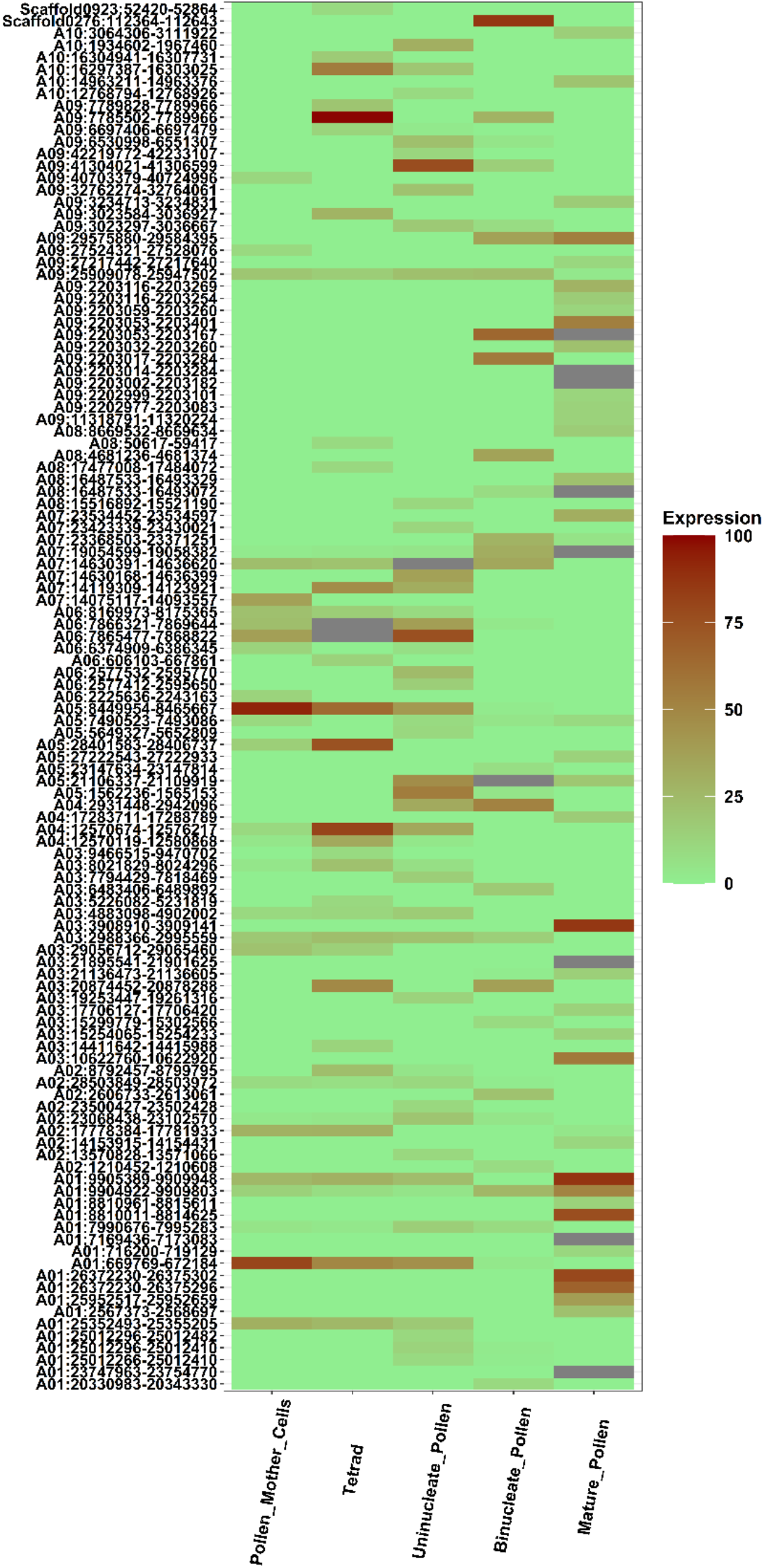
A heatmap showing the expression pattern of the top 60 (α ≥ 0.001) differential expressed circRNAs during pollen development in *B. rapa*. Gray tiles represent the expression of more than 100.

### Functional Annotation Analysis of CircRNAs Parent Genes

To explore the potential functions of circRNAs during pollen development, we performed GO enrichment analysis on the parental genes of all the differential expressed circRNAs. The results showed that the parent genes belonged to three categories: biological process, molecular function, and cellular component (TableS5). For the biological process, the circRNA-host genes were enriched in a variety of metabolic, catabolic, and cell processes such as protein folding, glucan catabolic process, regulation of phosphorus metabolism, meiotic DNA double-strand break formation process, meiotic cell cycle process, meiosis I cell cycle process and ncRNA metabolic process. In the molecular function, the enriched GO terms included GTPase activity, calmodulin-binding, catalytic activity acting on RNA, kinase regulator activity, and metal ion binding. In the cellular component category, only four terms, cytoplasm, endoplasmic reticulum, microtubule-associated complex, and katanin complex, were enriched (Figure 6).

**Figure 6.**
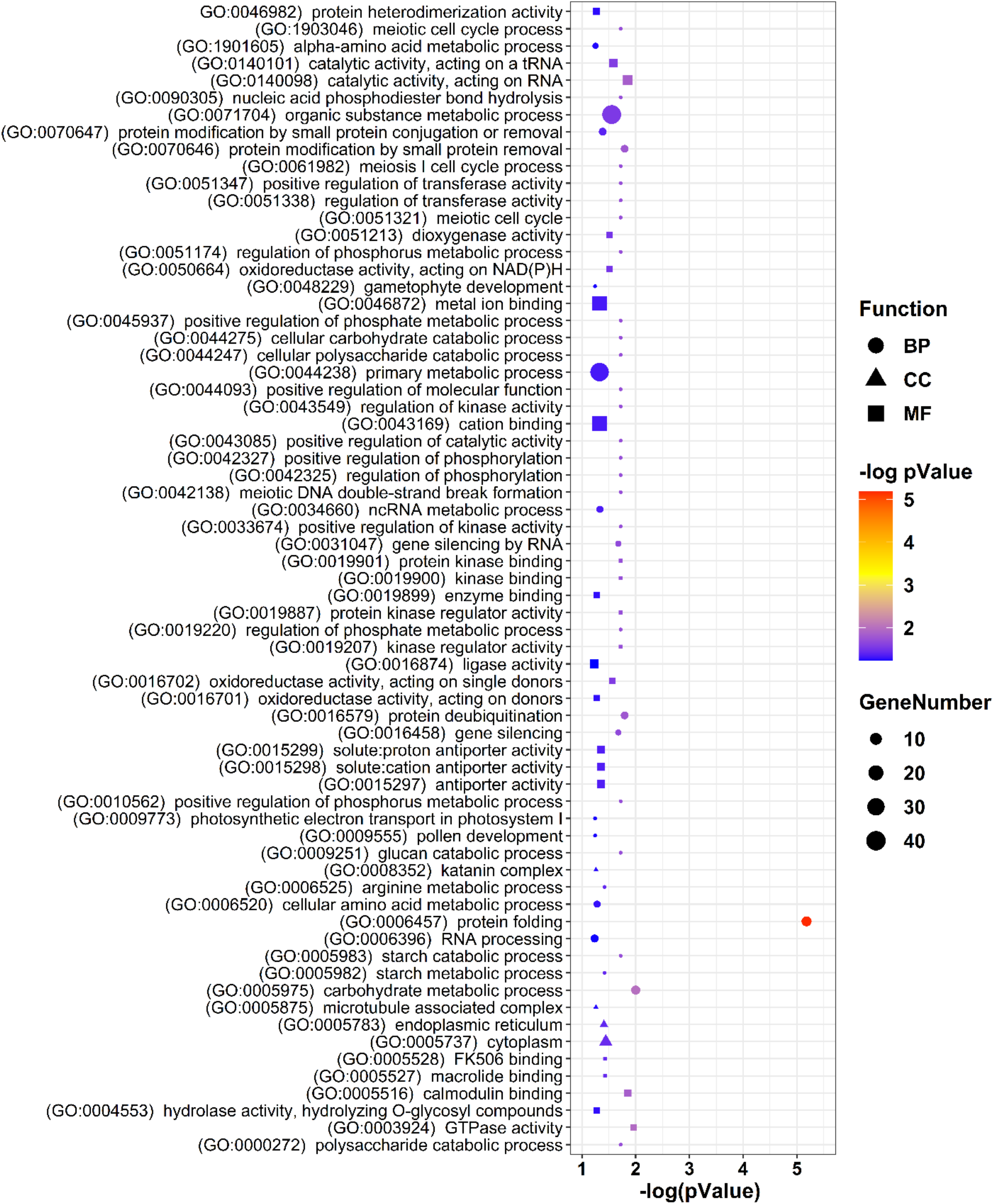
GO enrichment analysis of the host genes of differentially expressed circRNAs during pollen development in *B. rapa*.

To further investigate the function of circRNA host genes, Encyclopedia of Genes and Genomes (KEGG) pathway enrichment analysis was conducted. The results revealed enrichment of 23 significant pathways (TableS6), including Cyanoamino acid metabolism, protein processing in the endoplasmic reticulum, peroxisome, and aminoacyl-tRNA biosynthesis, among others (Figure 7).

**Figure 7.**
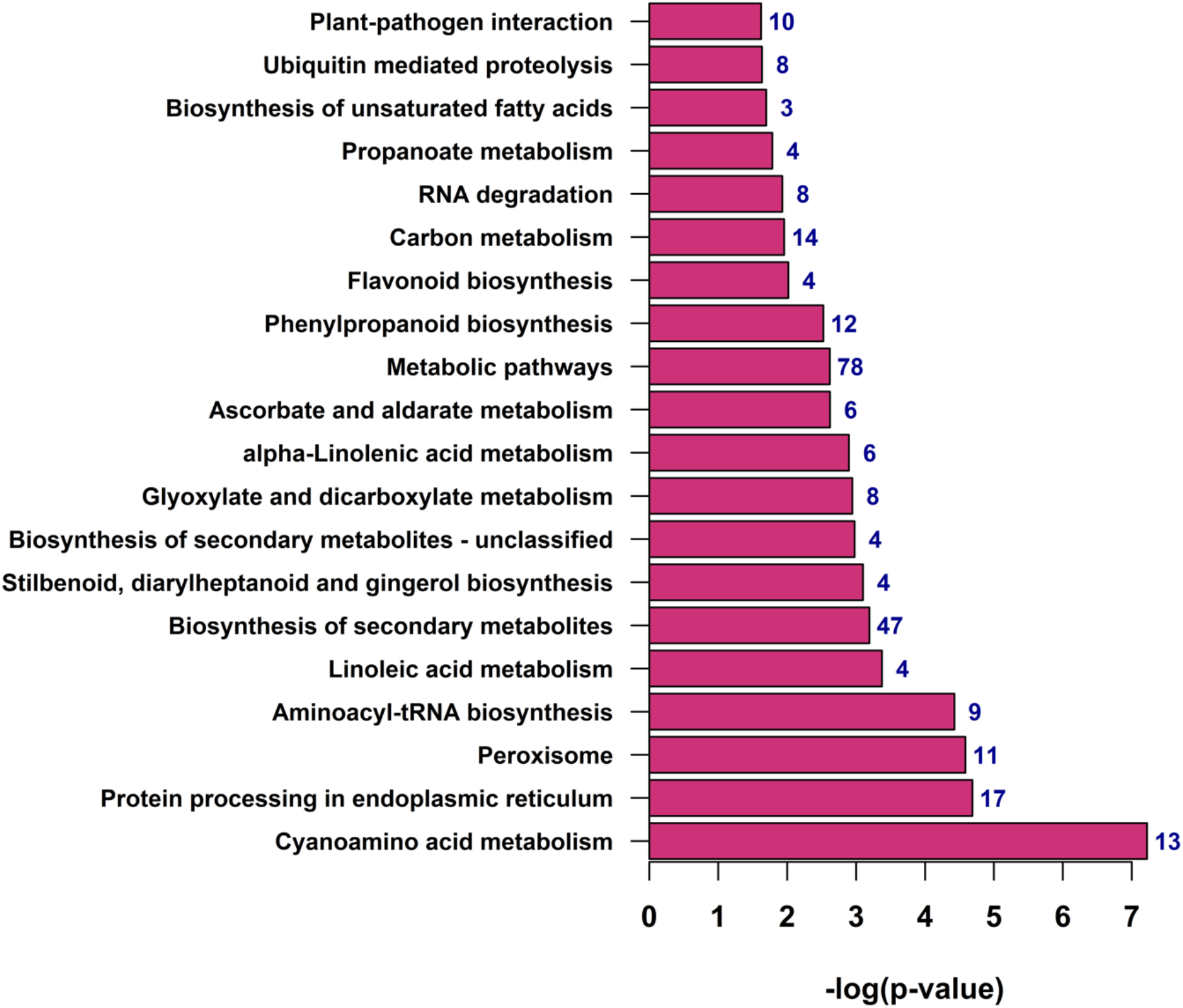
KEGG enrichment analysis of the host genes of differentially expressed circRNAs during pollen development in *B. rapa*. The number on the bars shows the number of genes enriched in the corresponding pathways.

### Prediction of miRNA target sites in circRNAs

To further evaluate the function of circRNAs in the post-transcriptional regulation of genes in pollen development, we predicted the binding sites of miRNAs in the sequences of differential expressed circRNAs (TableS7). The results showed that 88 circRNAs contained putative miRNA-binding sites for 73 miRNAs. Of these 88 circRNAs, 49 (55.68%) had only one miRNA binding site, followed by 22 circRNAs (25%) with two miRNA binding sites and the remaining circRNAs (19.32%) contained binding sites for three to nine miRNAs (Figure 8). For example, circRNA A06:606103-667861 and A09:10484746-10513688 had binding sites for eight and nine miRNAs, respectively. MiRNAs could be targeted by one or more circRNAs as well. Among 74 identified miRNAs, 27 (36.49%) were targeted by a single circRNA, 24 miRNAs (32.43%) were targeted by two circRNAs, and the remaining miRNAs (31.08%) were targeted by three to twelve circRNAs (Figure 8). For instance, bra-miR9569-5p, bra-miR5716, and bra-miR9563a-3p were targeted by seven, ten, and twelve circRNAs, respectively.

**Figure 8.**
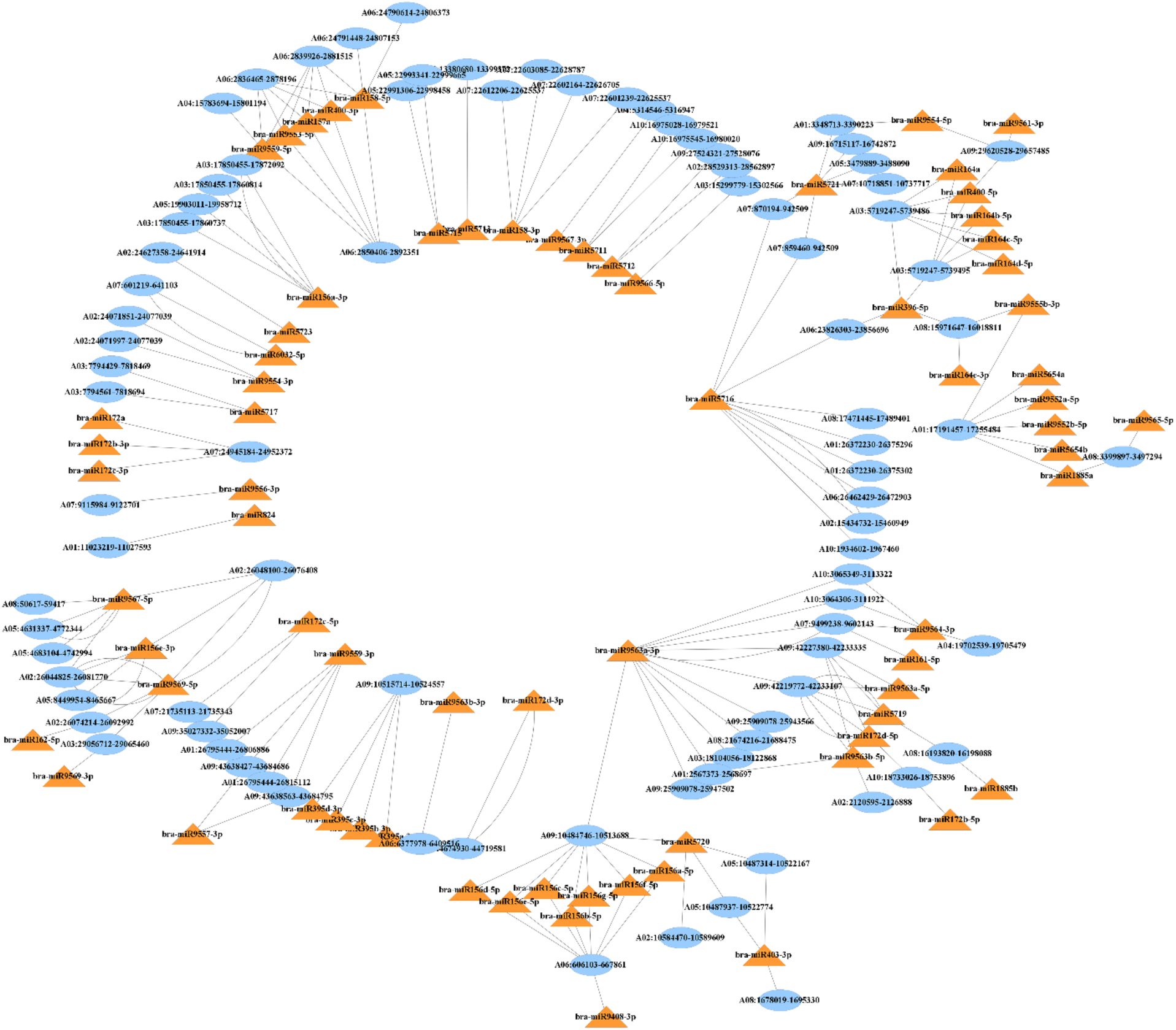
CircRNA-miRNA interaction network for differentially expressed circRNAs during pollen development in *B. rapa*. Blue ovals: circRNA IDs, Orange triangles: miRNA IDs.

### Validation of circRNAs using divergent primers

To confirm the accuracy of the identified circRNAs, we selected 15 circRNAs for experimental validation using PCR and Sanger sequencing. Using a set of divergent primers designed for each circRNA, we successfully amplified nine circRNAs using nine pairs of divergent primers. Then, all the amplified PCR products were further validated by sequencing to confirm the presence of the back-spliced junctions (Figure 9, Supplementary 2). Our validated circRNAs are composed of one to three exons, except for circRNA A03:5712452-5713124 which has an intron in its conjunction site.

**Figure 9.**
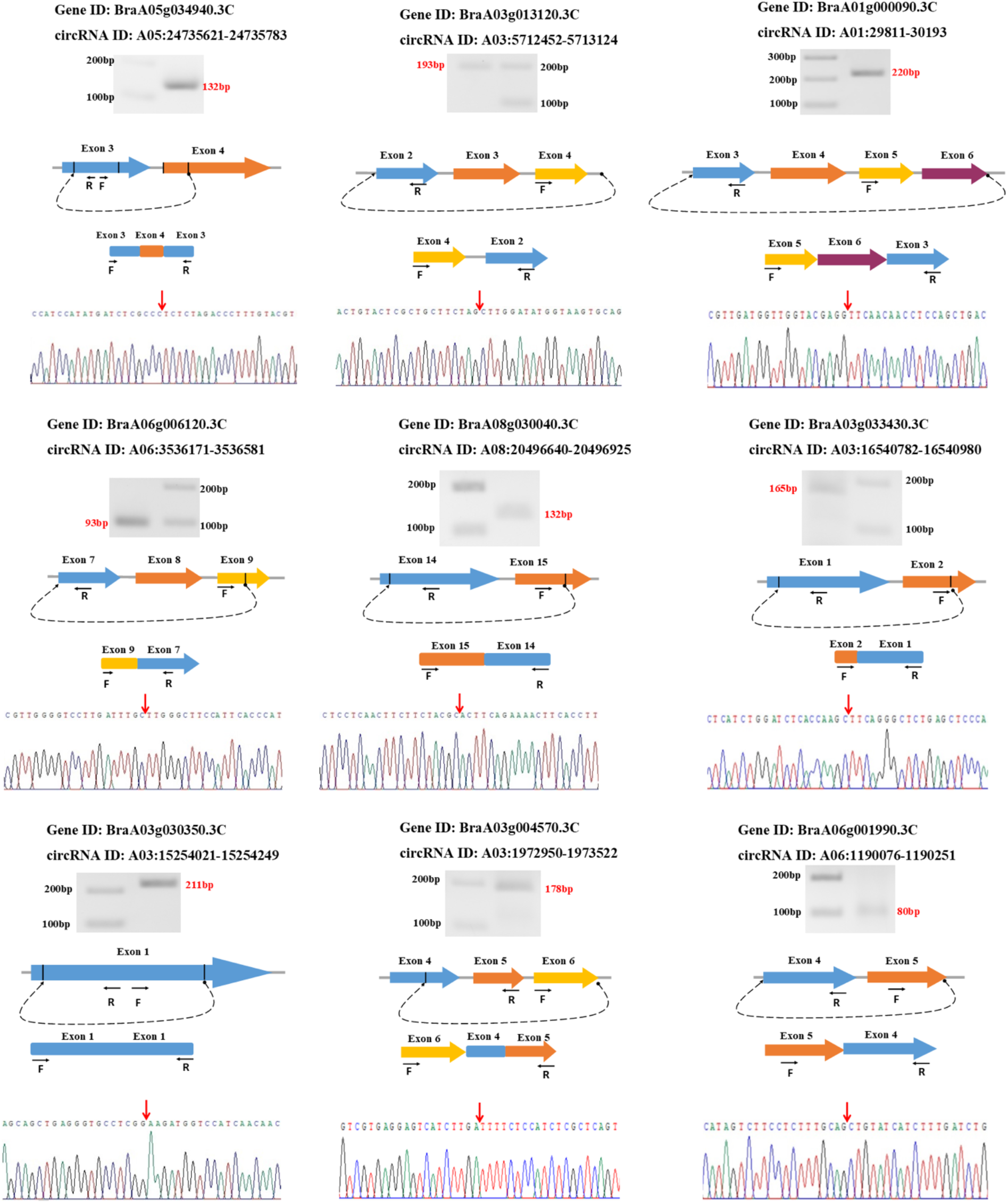
CircRNA validation using divergent primers and Sanger sequencing. From the top: the name of circRNA parent gene, circRNA ID, the amplified band using divergent primers and cDNA, schematic representation of the genomic region of selected circRNAs and their amplified junction, dashed curved arrows represent head-to-tail back-splicing points, the two sets of arrows indicate the position of divergent primers on corresponding exons, conjunction validation using Sanger sequencing, red arrows show the position of the junction point.

## Discussion

There is increasing evidence that circular RNAs are not simply rare splicing events but iare highly regulated molecules with unique selected sequences from different genomic regions (Wilusz 2018). CircRNAs can carry out important biological functions in a cell-type, tissue-, or developmental-stage-specific manner (Ebbesen et al. 2016). Although most of the circRNAs express at a low level compared with their cognate mRNAs, they can play important roles because they are naturally resistant to diverse exonucleases, and their half-life is much longer than mRNAs (Lasda and Parker 2014; Wilusz 2018). To date, thousands of distinct circRNAs have been found in various taxa, including different species of plants (Glažar et al. 2014; Chu et al. 2017). Various studies have suggested that plant circRNAs are closely associated with growth and development (Liu et al. 2017; Yang et al. 2020; Cheng et al. 2018). In the present study, we investigated the expression and potential function of circRNAs during pollen development in *B. rapa*. In total, we identified 1180 circRNAs in all five stages of pollen development, and comparing with previous reports; this number is similar to other plant species such as *Arabidopsis* (970) (Sun et al. 2016), *Gossypium arboretum* (1041), and *Gossypium raimondii* (1,478) (Zhao et al. 2017). We found that during pollen development, circRNAs originated from different genomic regions, but most of our identified circRNAs (87.63%) were generated from exonic regions, which is consistent with previous studies on many plant species such as *Arabidopsis* (85.1%) (Chen et al. 2017), Chinese cabbage (70.49%) (Wang et al. 2019b) and *Brassica campestris* (63.13%) (Liang et al. 2019). Most of the genes can produce only one isoform of circRNAs; however, multiple isoforms of circRNAs produced through alternative back-splicing from a single gene have also been observed (Zhang et al. 2019; Lu et al. 2015; Lv et al. 2020). Here we found that most of the circRNA parent genes (about 83%) generate only one circRNA isoform, with the remaining 17% of the genes able to produce two or more circRNA isoforms.

The expression of circRNAs changes during different stages of growth and development. For instance, by analyzing the expression profile of circRNAs in various plant species, it was revealed that circRNAs are closely involved in leaf growth in *Arabidopsis* (Liu et al. 2017), fruit ripening in pepper (Yang et al. 2020), and pollen development in *B. campestris* (Liang et al. 2019), rice (Wang et al. 2019c) and soybean (Chen et al. 2018). Herein, our differential expression analysis revealed the stage-specific expression pattern of circRNAs changes during pollen development. We found 130 differentially expressed circRNAs while pollen mother cells develop to tetrad, and 191 differentially expressed circRNAs while binucleate pollen develops to mature pollen. We noted that the most common circRNAs share between two closely related stages. For example, 272 shared differentially expressed circRNAs when tetrad develops to microspores and then to binucleate pollens. These findings indicate that circRNAs expression is related to different pollen developmental stages, and they have the potential to play important roles during pollen development in *B. rapa*.

Previous studies showed that circRNAs could regulate various processes such as chromatin structure, gene expression, translation, and even cell division (Liu et al. 2020; Li et al. 2015; Conn et al. 2017). For example, centromeric-retrotransposon-derived circRNAs in maize can bind to centromeres via R-loops to promote chromatin looping in centromere regions (Liu et al. 2020). Exon-intron-containing circRNAs in the human cells can increase the transcription of their parental genes by interacting with U1 small nuclear ribonucleoproteins at the promoters (Li et al. 2015), or in *Arabidopsis*, circRNAs can form a strong RNA-DNA hybrid resulting in transcriptional pausing (Conn et al. 2017). CircRNAs also can form circRNA-protein structures and affect translation and cell cycle regulation. One such example is CircPABPN1 which can suppress HuR from binding to its cognate mRNA, resulting in reduce translation of PABPN1 mRNA (Abdelmohsen et al. 2017). Another example is circFOXO3 which has multiple binding sites for cell cycle regulating proteins such as p53, CDK-2, and p21. It was reported that circFOXO3, p21, and CDK-2 could form a ternary structure and inhibit the CDK-2/Cyclin-E complex activation, which is necessary for G1 to S phase transition (Du et al. 2016). During pollen development, several mitotic and meiosis cell divisions occur, which involve the expression of a few thousand genes (Rutley and Twell 2015; Singh and Bhalla 2007). Studies showed that during pollen development, the tapetum endoplasmic reticulum was highly involved in biosynthesis, folding, and secreting proteins (Fragkostefanakis et al. 2016; Singh et al. 2021). For pollen wall formation, lipidic components and polysaccharides are required as tapetum secretes lipid components onto the pollen surface (Jiang et al. 2013; Sharma et al. 2015), or β-glucosidase, which is involved in the regulation of polysaccharide metabolism, downregulated in the sterile floral buds of *B. rapa* (Dong et al. 2013). As circRNAs function may be related to their parent gene function, we annotated the biological roles of circRNAs parent genes using GO and KEGG analysis better to understand the potential function of circRNAs in pollen development. We noted that circRNA parent genes could be involved in various important processes and functions related to pollen development such as protein synthesis and fate, cell cycle and DNA processing, kinase and phosphorus activities, polysaccharide metabolism, gene silencing, RNA processing and antiporter activities. KEGG analysis showed that the circRNAs parent genes are involved in 23 pathways, mainly amino acid metabolism and protein processing, lipid biosynthesis and metabolism, metabolic pathways, carbon metabolism, and RNA degradation. Based on our results and previous findings, we can assume that circRNAs might play vital functions during pollen development in *B. rapa*.

Small RNAs such as miRNAs can regulate gene expression through mRNA cleavage, translational repression, or gene silencing via miRNA-directed DNA methylation (Waititu et al. 2020). Studies on animal and human cells have reported that circRNAs could bind to miRNAs and sequester them from their target mRNAs to regulate the gene expression at the post-transcriptional level (Li et al. 2018). For example, it has been shown that *circSry* associated with testis development in mice, contains 16 binding sites for miR-138 (Hansen et al. 2013), or *CDR1as*, which is a highly expressed circRNA in mammalian brains, contains more than 70 binding sites for miR-7. Studies showed that *CDR1as* regulate gene expression by acting as miR-7 storage (sponge) and ensuring the release of an appropriate amount of miR-7 to the target mRNAs (Kleaveland et al. 2018; Piwecka et al. 2017; Xiao et al. 2020). Prediction of circRNA-miRNA networks in plant species revealed that plant circRNAs, compared with animals and humans, have fewer interactions with miRNAs suggesting that the main function of plant circRNAs might not be a miRNA decoy (Chu et al. 2020). Nevertheless, several studies proposed that plant circRNAs could target miRNAs by acting as competing endogenous RNAs in post-transcriptional regulation of the genes (Frydrych Capelari et al. 2019; Liang et al. 2019; Liu et al. 2019; Wang et al. 2019c; Chen et al. 2018). For example, circRNAs were proposed to function as miRNAs sponges in flower development in *Arabidopsis* (Frydrych Capelari et al. 2019), anther development in *B. campestris* (Liang et al. 2019), and pollen development in rice and soybean (Wang et al. 2019c; Chen et al. 2018). Even a database named “GreenCircRNA” collected all the identified circRNAs in different plant species and predicted circRNA-miRNAs interactions for each species (Zhang et al. 2020a). In the present study, we found 88 circRNAs with putative miRNA binding sites. A few of them had multiple binding sites for the same or different miRNAs suggesting that circRNAs could interact with miRNAs or act as miRNA sponges to regulate pollen development. We also noted some well-known and important miRNAs in our predicted network, such as miR156, miR157, miR158, miR161, miR162, miR164, miR172, miR395, and miR396. These miRNAs proved to be involved in many biological and developmental processes such as vegetative growth, flowering, fertility, and fruit ripening (Waititu et al. 2020; Luján-Soto and Dinkova 2021; Wu 2013; Li and Zhang 2016). For example, we found nine circRNAs with binding sites for miR156, seven circRNA with binding sites for miR172, four circRNAs with binding sites for miR396, and three circRNAs with binding sites for miR164, which all are known to regulate flower development in *Arabidopsis* (Wang et al. 2009; Jung et al. 2014), barley (Nair et al. 2010), tomato (Cao et al. 2016), *Brassica napus* (Wang et al. 2019a), and strawberry (Zheng et al. 2019). In addition, we found nine circRNAs with binding sites for miR158 that were previously reported as key regulatory miRNA during pollen development in *Brassica campestris* (Ma et al. 2017). Accordingly, our results suggest that circRNA could act as ceRNAs during pollen development in *B. rapa*.

## Conclusion

In conclusion, by studying the profile of circRNA expression during pollen development in *B. rapa*, we identified 1180 circRNAs, of which 367 circRNAs showed stage-specific expression. Functional characterization of circRNAs host genes revealed that circRNAs mainly were related to biological and molecular processes in pollen development such as mitosis and meiosis cell cycles, protein biosynthesis, protein modification, and polysaccharide processes. Moreover, 88 circRNAs were found to contain miRNA binding sites suggesting the role of circRNAs in gene expression as post-transcriptional regulatory elements. Our study revealed the potential functions of circRNAs during pollen development and paved the way for further experiments on studying the molecular mechanism of these new regulatory RNA molecules.

## Supplementary Materials

The following are available online at (www.mdpi.com/xxx/s1), Table S1: All the circRNAs found using three algorithms in five pollen developmental stages, Table S2: List of 1180 unique circRNAs including annotation, Table S3: The expression pattern of circRNAs during pollen developmental stages, Table S4: Normalized expression of circRNAs in each pollen developmental stage, Table S5: GO annotation of circRNAs’ parental genes, Table S6: KEGG pathway enrichment of host genes of differentially expressed circRNAs, Table S7: Predicted miRNA-circRNA interactions. Supplementary Doc.: The nucleotide sequence of validated circular RNAs’ junctions.

## Author Contributions

Conceptualization: S.B., M.B.S. and P.L.B.; validation: S.B.; formal analysis: S.B.; Resources: S.B., M.B.S. and P.L.B.; Original draft preparation: S.B.; visualization: S.B.; review and editing: M.B.S. and P.L.B.; Supervision: M.B.S. and P.L.B. All authors have read and agreed to the published version of the manuscript.

## Funding

This research was funded by the University of Melbourne Research Scholarship.

## Data Availability Statements

All data generated in this study are available in the article and its supplementary files.

## Acknowledgments

This research was supported by Melbourne Bioinformatics at the University of Melbourne, project punim1093.

## Institutional Review Board Statement

“Not applicable”

## Informed Consent Statement

“Not applicable”

## Conflicts of Interest

The authors declare no conflict of interest.

## Notes

### Competing Interest Statement

The authors have declared no competing interest.

